# A network of transcriptomic signatures identifies novel comorbidity mechanisms between schizophrenia and somatic disorders

**DOI:** 10.1101/2023.10.02.560428

**Authors:** Youcheng Zhang, Vinay S. Bharadhwaj, Alpha T. Kodamullil, Carl Herrmann

## Abstract

The clinical burden of mental illness, in particular schizophrenia and bipolar disorder, are driven by frequent chronic courses and increased mortality, as well as the risk for comorbid conditions such as cardiovascular disease and type 2 diabetes. Evidence suggests an overlap of molecular pathways between psychotic disorders and somatic comorbidities. In this study, we developed a computational framework to perform comorbidity modeling via an improved integrative unsupervised machine learning approach based on multi-rank non-negative matrix factorization (mrNMF). Using this procedure, we extracted molecular signatures potentially explaining shared comorbidity mechanisms. For this, 27 case-control microarray transcriptomic datasets across multiple tissues were collected, covering three main categories of conditions including psychotic disorders, cardiovascular diseases and type II diabetes. We addressed the limitation of normal NMF for parameter selection by introducing multi-rank ensembled NMF to identify signatures under various hierarchical levels simultaneously. Analysis of comorbidity signature pairs was performed to identify several potential mechanisms including 1) dysfunction of endothelial systems; 2) induction of hypoxia and oxidative stress; 3) dysregulation of neural transmission GABAergic system associated with neuroendocrine function (e.g. insulin secretion); 4) activation of inflammatory response auxiliarily interconnecting blood-brain systems, oxidative response and GABAergic neuro-action. Overall, we proposed a general cross-cohorts computing workflow for investigating the comorbid pattern across multiple symptoms, applied it to the real-data comorbidity study on schizophrenia, and further discussed the potential for future application of the approach.

## Introduction

Mental illnesses, in particular schizophrenia and bipolar disorder, are increasingly raising public health awareness globally. The clinical burden is not only driven by frequent chronic courses and increased mortality but also by the risk for somatic comorbid conditions such as cardiovascular disease ^1^ and type II diabetes ^2^. People with mental illnesses often show an increased susceptibility to somatic illnesses. In fact, the comorbidity between psychotic diseases and somatic illnesses is mutual ^3^, with studies showing a higher risk of developing a psychiatric disorder for persons affected by certain somatic diseases ^1,3^. Conversely, the risk of somatic disease is approximately twice as high for persons with severe mental illnesses than for people without mental disorders ^3^. Studies on genetic susceptibility have suggested a potential overlap of molecular mechanisms between psychotic disorders and somatic comorbidities ^4,5^, such as shared gene risk loci that were demonstrated in several genotype studies ^6^, biomarkers that were dysregulated across psychosis and comorbid diseases in transcriptomic analysis ^7,8^, and shared molecular pathways that were identified in integrated analysis ^7^.

To explore shared molecular mechanisms between somatic comorbidities and psychotic disorders, previous studies have been conducted with different approaches. For instance, genetic overlap analyses were performed on genome-wide summary statistics from two large-scale GWAS of schizophrenia and type II diabetes to discover the shared association between schizophrenia and type II diabetes ^6^. Common differential expressed genes (DEGs) were analyzed as the comorbid genes across schizophrenia and type II diabetes and further identified the enriched gene ontology as well as transcription factors with these DEGs ^9,10^. Similarly, a set of susceptibility genes for schizophrenia and type II diabetes were retrieved respectively to identify the significant pathways crossing among two syndromes ^10^. Mendelian randomization analysis was performed to establish the causal linkage of genetic variants associated with type II diabetes with the risk of schizophrenia using an inverse-variance weighted meta-analysis ^11^. Multi-trait colocalization, which suggested a common genetic etiology between schizophrenia and cardiometabolic traits, was estimated with linkage disequilibrium score regression (LDSC), genetic covariance analyzer (GNOVA), and heritability estimation from summary statistics (ρ-HESS) ^12^, whereas polygenic risk scores were included additionally to identify the potential shared genetic variants and inferred the pathways involved ^13^. The genetic-pleiotropy-informed method was introduced for improving gene discovery with the use of GWAS summary-statistics data to identify additional loci associated with schizophrenia, connecting various cardiovascular disease risk factors ^14^. Molecular correlations and bi-directional Mendelian randomization (MR) were computed to assess the causality of an increased risk of cardiovascular disease in schizophrenia patients and to explore the underlying mediating factors ^1^. Moreover, metagenes and molecular pattern discovery using independent component analysis (ICA) ^15^ in independent transcriptomes provided us new insights to study the molecular comorbidity^16–18^.

However, due to the limited knowledge as well as the lack of systematic analysis on the comorbidity in psychotic disorders, the underlying mechanism remains to be explored. Therefore, in this study, we performed a systematic comorbidity analysis by developing a computational framework modeling leveraging a large-scale transcriptomic dataset via an improved integrative unsupervised machine learning approach based on multi-rank non-negative matrix factorization (mrNMF). Using this procedure, we extracted molecular signatures potentially explaining shared comorbid mechanisms. For this, 27 case-control microarray cohorts across multiple tissues were collected, covering three main categories of conditions including psychotic disorders, cardiovascular diseases, and type II diabetes. We addressed the limitation of normal NMF for parameter selection (e.g. undetermined number of ranks) by introducing multi-rank ensembled NMF to identify signatures under various hierarchical levels simultaneously. A reciprocal best-hit scoring matrix was computed to integrate signatures across different cohorts and to generate the disease signature graph while controlling confounding effects. Biological representation and gene analysis based on signature pairs of the graph was performed to identify comorbid processes and marker genes. GWAS mapping with the risk variants of each trait was implemented to validate the key genes in our analysis. Finally, we queried an independently curated knowledge graph to reveal the underlying relationships among shared molecular factors and the co-occurrence of these biological processes, further helping bridge the roadmap of the mechanism of the comorbidity.

## Method

### Datasets

27 case-control cohorts with 1163 individuals in total (633 patients and 530 healthy participants) were retrieved from Gene Expression Omnibus (GEO) repository, covering three main categories of conditions including psychotic disorders, cardiovascular diseases, and type II diabetes (See Supplementary Table S1). To reduce the system biases induced by different platforms, we only selected datasets using platform GPL570 [HG-U133_Plus_2] Affymetrix Human Genome U133 Plus 2.0 Array. Within the three main categories of conditions, various subtypes were collected: in the psychotic disorders category (PSY), schizophrenia (scz), bipolar disorders (bd), and major depression (mdd) were included, whereas for cardiovascular diseases (CVD), acute coronary syndrome (acs), coronary artery disease (cad), acute myocardial infarction (ami), peripheral arterial disease (pad), hypertension (hyp), and cardiomyopathy (cdm) were included. Categories are indicated in upper-case (PSY, CVD, T2D) while individual diseases are in lower-case. Each cohort involved individual samples from only one tissue, including brain, blood, islet, liver, etc. Other variables such as gender, age, BMI, etc, available in cohorts were extracted and stored in the metadata for further processing. Details were summarized in supplementary Table 1.

### Data preprocessing

All the raw expression microarray data were retrieved from GEO as .CEL files. Frozen robust multiarray analysis (fRMA) ^19,20^ was performed to preprocess raw data into expression data. RMA background correction, quantile normalization and robust weighted average summarization were applied to the fRMA preprocessing pipeline setting. Sample and outliers with missing values were removed, along with their corresponding expression data. Probes were mapped to gene symbols using biomaRt ^21^ package. Finally, we performed rowmean normalization (setting the mean value of each gene to 1).

### Multi-rank non-negative matrix factorization (mrNMF)

Multi-rank non-negative matrix factorization (mrNMF) is an improved version of non-negative matrix factorization (NMF). To discover the hidden structure in high-dimensional expression data, multi-rank non-negative matrix factorization decomposes the data matrix with genes in rows and samples in columns into two component matrices, gene-rank matrix (W) and rank-sample matrix (H). Here, the rank refers to the “signature” (column of the W matrix) which is a weighted genes vector, in which the weights are the exposure. The value of the rank k is a tunable parameter, referring to the number of signatures. However, instead of selecting a single value for k, mrNMF was applied with a series of rank values from k = 2 to k = 20. Then all the resulting gene-signature W matrices were concatenated into one large W matrix while the signature-sample H matrices were concatenated into one large H matrix The loss function for each round of mrNMF with specific rank k is provided below.

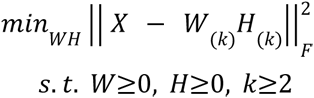

The ensembling was implemented by augmenting all the W matrices and H matrices respectively, where the concatenation is implemented by column for W matrices and by row for H matrices.

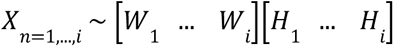

We call “disease signature” the columns of the W matrix, representing the positive contributions of all genes to this signature.

### Identification of diagnosis-associated signatures

To identify the signatures that were significantly associated with the disease phenotype, a linear model was applied with values of each of the signatures in rank-sample H matrices as independent variables, group variable (case or control) as dependent variable and gender, age, and other available variables as covariates. A linear model with a t-test was performed to identify the signatures associated with the biological effect (P-value < 0.05). The corresponding column in the gene-rank W matrix and row in the rank-sample H matrix was kept while removing the non-significant signatures.

### Similarity measurement of the signatures

In our graph approach, links between signatures were defined by measuring the similarity between two signatures. We assume that only a few genes significantly contribute to the signature, whereas the weight of most genes corresponds to random noise. Hence, instead of using correlation over all genes, we computed the Jaccard similarity coefficients using the top N (N=1000) genes (based on the exposure values) between each pair of signatures from two different cohorts in gene-rank W matrices (Supplementary Figure S1). Here we also compared the differences by using different values of N (Supplementary Figure S2).

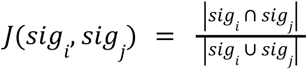

Finally, we define *g_N_* (*i*, *j*) as the list of shared genes among the top N genes in signatures i and j.

### Reciprocal best hit scoring

The principle of Reciprocal best hit (RBH) was previously introduced in comparative genomics to determine orthologous genes between pairs of species. Reciprocal best hit scoring is a metric to obtain the best concordance of pairs of signatures from different individual cohorts. We used Jaccard similarity coefficients (see previous section) as the scoring criterion for signature pairs. For each signature A obtained from one dataset, we determined the most similar signature B from all other datasets and verified if signature A was also the most similar signature to signature B. Hence, we only considered the similarities between signatures extracted from different datasets, and not between those extracted within one dataset. In that case, A and B are called RBH. We split the signatures into two groups, 1) positive signature and 2) negative signature, based on the t-value computed in the previous step “Identification of diagnosis-associated signatures” to represent the orientation of the signatures towards the diagnosis group variable. The RBH scoring was computed only within the signatures in the same group. All the best hits signature pairs were connected by an edge in the signature graph.

### Signature graph construction

The signature graph was built with input from the reciprocal best hit scoring step. In the graph, each node represented one individual signature from one cohort. The attribute of the nodes contains information including tissue, disease category, subtype, and diagnosis association. The edges represent RBH relationships, with scoring values as the edge weights.

### Permutation test

To evaluate the empirical significance of the edges, a permutation procedure was implemented. We performed randomization control to identify the signature pairs that existed more than expected under a randomized case. The exposure values in gene-rank W matrices obtained in the mrNMF decomposition step were shuffled. Similarity measurements of the signatures, reciprocal best hit scoring, and graph construction were then conducted subsequently. This procedure was repeated 20 times and the permutation test was computed to observe the occurrence of specific edges in the actual graph with unshuffled data on the permuted graph with shuffled data. If the probability of the occurrence of a specific edge is lower or equal to 1 out of 20 (P ≤ 0.05), this edge would be considered significant. Insignificant edges with P > 0.05 were removed.

### Comparability of the signatures across different cohorts

To make signatures extracted from different cohorts comparable, min-max scaling normalized the gene exposure of each signature to a min-max of [0, 1], considering both the minimum and maximum values of the signature.

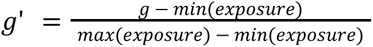

### Identification of different types of comorbid gene

We first defined a signature pair that links two different diseases as a comorbid signature pair (e.g. scz-bd, scz-t2d) whereas a signature pair that links the same disease is defined as a single disease signature pair (e.g. scz-scz, t2d-t2d). As described earlier, each signature pair has its corresponding gene list of top genes shared between the two individual signatures. We collected the union of all genes shared by single-disease signature pairs. For example, we define the union of all genes shared by all scz-scz signatures pairs, and likewise for the t2d-t2d and other single-disease pairs. We consider these genesets as representing genes involved in processes specific to a disease (Figure 3b & Supplementary Figure S1). For simplicity, we refer to these genes as, e.g., “schizophrenia-associated genes”. For the genes shared by a comorbid signature pair that links e.g. schizophrenia to diabetes (referred to as “comorbidity-associated genes”), we categorized the genes into four types including “schizophrenia-specific genes” (genes that are only found in the set of schizophrenia-associated genes), “diabetes genes” (genes that are only found among the diabetes-associated genes), “comorbid genes” (genes that are found in both schizophrenia and diabetes associated genes) and “specific genes that overlap with neither schizophrenia nor comorbidity genes” (genes that are not found in single disease signature pairs).

### Biological inference on signature pairs

Biological interpretation on the edges, in terms of enriched biological processes, was performed using the ClusterProfiler package ^22^ to perform enrichment analysis. For each edge, the overlapping gene list *g_N_* (*i*, *j*) obtained from the previous step was used as the input. Gene Ontology resource (GO) from MsigDB ^23,24^ was used. FDR correction was implemented to keep the significant GO terms (adjusted P-value < 0.05).

### Gene analysis on specific biological processes of signature pairs

Enriched genes in specific ontology terms (*acute inflammation*, *angiogenesis*, *oxidative stress,* and *GABAergic synapse*) were explored in detail. We first extracted the scz-t2d signature pairs that showed enrichment of one of these four ontology terms, meanwhile, we retrieved the enriched genes of these terms. For each gene and each signature pair, we obtained two exposure values, one from the schizophrenia signature, and the other from the comorbidity signature within the signature pairs. Genes with top 30 exposure value were ranked based on the geometric mean of the exposure values in the two individual signatures of the signature pair and visualized in a scatter plot with the exposure value for schizophrenia and the comorbid disease over the signature pairs as x and y, whereas we also annotated the types of genes retrieved from the previous section (*distinct, shared, schizophrenia specific, comorbid specific*).

### Mapping of the embedded genes in specific comorbid mechanism on the GWAS data

To validate the gene list identified in the scz-t2d comorbid signature pairs, we mapped the genes enriched in *acute inflammation*, *angiogenesis*, *oxidative process* and *GABAergic system* to the GWAS risk variants from schizophrenia and type 2 diabetes. 4988 associations in 142 studies with corresponding mapped gene information for schizophrenia and 5263 associations in 218 studies for type 2 diabetes were retrieved from GWAS catalog. The shared GWAS genes between schizophrenia and type 2 diabetes were treated as comorbid GWAS genes. The comorbid gene that we found in schizophrenia-t2d comorbid signature pairs were further summarized into different categories including “gwas_t2d”, “gwas_scz” and “gwas_shared”.

### Building the schizophrenia - type 2 diabetes comorbidity knowledge graph

We used a literature-curated knowledge-graph to build the connectivity paths from schizophrenia to type 2 diabetes and validate through an independent approach the specific processes identified from all the previous analyses in the study. Specific processes including *acute inflammation*, *angiogenesis*, *oxidative process,* and *GABAergic* were selected. We followed the procedure in the previous section to extract a list of query genes. The top 30 enriched genes with the highest exposure value of these processes were retrieved. With the selected processes and the corresponding high exposed genes as input, we fed them into a literature-curated knowledge graph database ^25^. The graph with maximal connectivity depth = 3 (connected node degree) involved the specific processes and genes above was generated. Based on the graph, we created and visualized the subgraph for each biological process we wanted to highlight, by starting with the biological process and searching for all the shortest paths to schizophrenia node and type 2 diabetes node respectively.

## Result

### Overview of the SigGraph computation framework

To investigate the biological dimensions shared between psychotic disorders and somatic comorbidities, we developed a graph-based computational framework to perform comorbidity modeling via an improved integrative unsupervised machine learning approach based on multi-rank non-negative matrix factorization (mrNMF). Compared with the normal non-negative matrix factorization (NMF), mrNMF is optimized in several aspects. For instance, the factorization rank *k* must be provided in standard NMF, where a high rank might identify spurious signatures with little biological meaning whereas a low rank results in only a few signatures that might not be able to separate different biological factors in detail. Instead, we perform a multirank decomposition between *kmin* and *kmax*. Therefore, mrNMF can be regarded as an ensemble strategy that considers multiple ranks simultaneously. Moreover, with increasing rank parameter *k*, the biological signal captured by the parent nodes will split into their descendant nodes (Figure 1), meaning that on different levels of *k*, signatures would capture different levels of the biological effects. With the multi-rank strategy, the model can identify signatures under different hierarchical levels coincidently. In the study, three large collections of transcriptomic datasets with 1163 individuals (633 patients and 530 healthy participants) were collected covering various psychotic disorders (schizophrenia, bipolar disorder, and major depressive disorder), type II diabetes and cardiovascular disease (coronary artery disease, peripheral artery disease, etc.) (Figure 1, Table S1). First, we applied mrNMF on each preprocessed transcriptomic dataset independently to decompose the data matrix into gene-signature *W* matrices and rank-sample *H* matrices (see Methods). The rank-sample *H* matrices were used to filter signatures (i.e. rows of the *H* matrix) significantly associated with diagnosis. Second, to infer correspondences between the gene signatures, we applied the reciprocal best-hit scoring as the similarity measurement metric on every pair of signatures obtained from different cohorts. As defined in the previous study ^16,18^, a reciprocal best hit is found when a pair of signatures, each in a different cohort, is identified with each other as the best scoring match. Reciprocal best hit scoring has already been shown effective in batch correction as well as the removal of confounding variables in the original high-dimensional space ^26^, by simply identifying the best matching pair of signatures from two different cohorts (Figure 1). However, estimating the relationship between two signature vectors which contain more than 20,000 genes using correlation might lead to spurious associations, as only the top-ranked genes should be expected to be associated with the biological condition. This is similar to the concept of the leading edge in gene set enrichment analysis (GSEA). Thus, rather than focusing on the global patterns of the gene exposure in each of these signatures, we focused on top exposed genes and computed Jaccard similarity between the top 1000 genes to measure the similarity of the signature pairs. We have verified that the results are not dependent on this parameter (Fig. S2). By doing so, we constructed the graph with edges weighted by Jaccard similarity between every best-matching signature pair. Third, to better understand the molecular origin of comorbidity, we performed an analysis on the graph components. Our hypothesis was that, when two connected signatures come from datasets corresponding to different diseases, the shared genes represent common molecular processes found in both diseases. Thus, the comorbidity information can be extracted from the signature pair (edge) level. However, some of the signature pairs might exist by chance rather than the real biological effect. To avoid this, we performed a permutation test by randomly constructing multiple graphs as controls and retained the signature pairs that occurred more than expected in the situation under randomization (see Methods). Thus, the signature pairs with true molecular effect tended to stay in the graph whereas the spurious pairs were removed. To identify the mechanism behind this, enriched biological processes were identified. Various types of comorbid genes were categorized and analyzed as well. Finally, to validate the results, GWAS data was used to validate the identified shared genes using genetic data.

**Figure 1.**
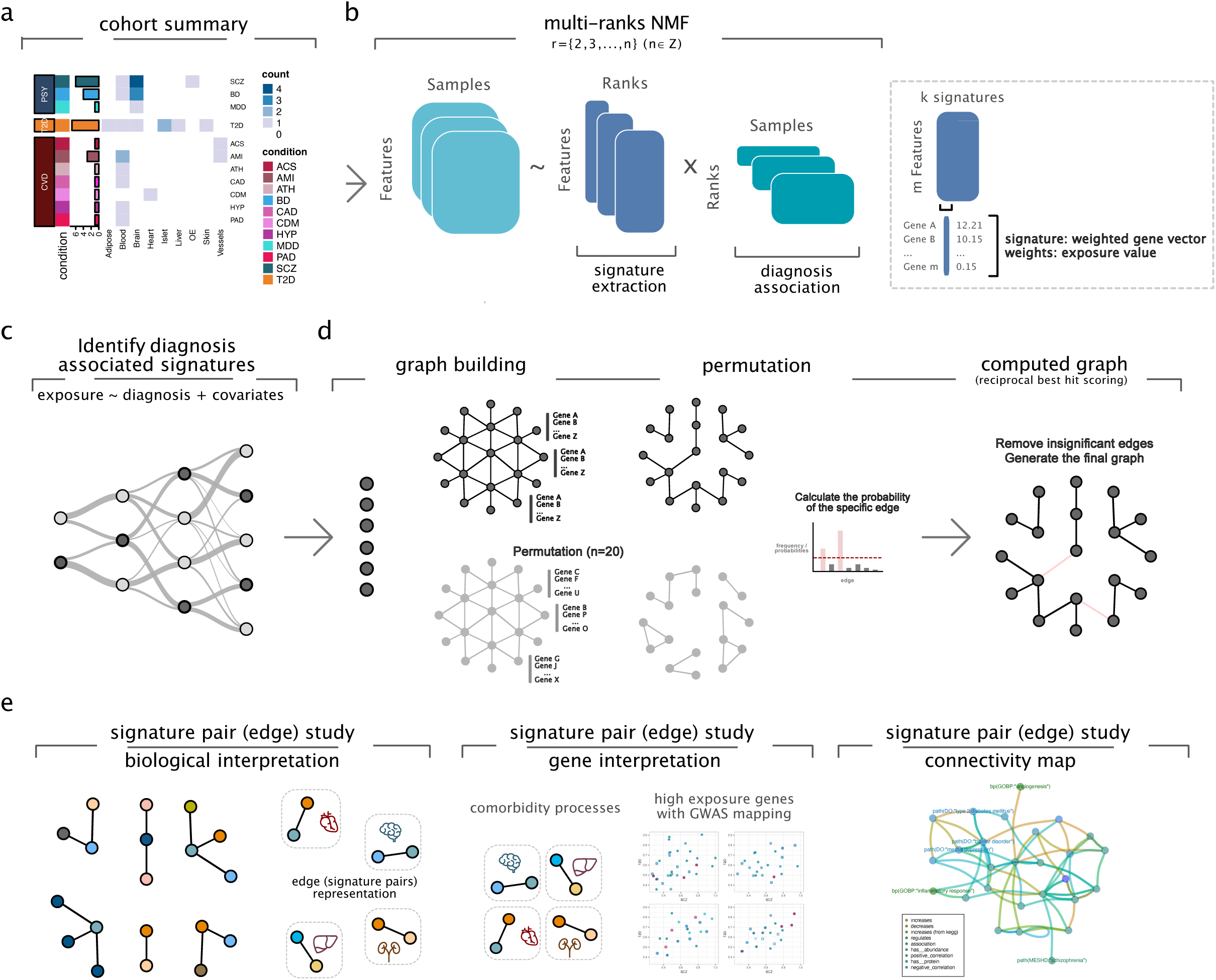
Overview of the signature graph computational framework based on multi-rank non-negative matrix factorization. **a** Descriptive summary of the transcriptomic cohorts, the number of cohorts by disease and tissue, scz (schizophrenia), bd (bipolar disorders), mdd (major depressive disorders), t2d (type 2 diabetes), acs (acute coronary syndrome), ami (acute myocardial infarction), ath (Atherosclerosis), cad (coronary artery disease), cdm (dilated cardiomyopathy), hyp (hypertension), pad (peripheral artery disease). **b** Algorithm of multi-rank non-negative matrix factorization. **c** Identification of diagnosis-associated signatures. **d** Graph building by reciprocal best hit matrices and permutation test. **e** Comorbidity analysis based on signature pairs including biological interpretation, high exposed gene identification and connectivity map establishment.

### Description of the derived signature graph

Our signature graph contains 419 signatures (represented as nodes) and 305 signature pairs (represented as edges) (Fig. 2a). These signatures belong to three different disease classes (PSY, T2D and CVD), 9 tissues and 58 types of edges (scz-scz, scz-t2d, bd-t2d, etc). To standardize the notation, all the notations referring to disease classes (e.g. PSY, T2D, CVD) are in uppercase while those for specific diseases (scz, t2d, bd, ami, etc.) are in lowercase. We also extracted a subgraph containing only schizophrenia and type 2 diabetes signatures (Fig. 2a). The edges between individual signatures suggest shared biological processes leading to similar transcriptomic outcomes. Hence, information about shared molecular effects can be derived from a detailed analysis of signature pairs. Therefore, to explore the graph in detail, we summarized all the signature pairs (Fig. 2b). We categorized four different groups of signature pairs, namely *single* (signatures from the same disease, e.g. scz-scz), *intra-class comorbid* (signatures from the same disease class e.g. scz-bd), *inter-class comorbid* (signatures from distinct classes e.g. scz-t2d) and *somatic* (signatures from non-psychotic diseases). Among all signature pairs falling into the inter-class comorbid category, those connecting a psychiatric disorder and a somatic disease, in particular, schizophrenia - type 2 diabetes (scz-t2d), were observed most frequently. We questioned whether the tissue of origin represented a main confounder of the signature matching, in the sense that signatures derived from the same tissues would be associated more frequently. However, not only did the edges connect signatures from different diseases, they also frequently connected signatures obtained from different tissues. Therefore, to examine the tissue effect, we also summarized the number of cross-tissue signature pairs (Fig 2b). It appears that signature pairs that linked individual signatures from different disease cohorts across different tissues were predominant, suggesting that tissue effect is not a main driver of signature clustering (Fig. 2b). Furthermore, within the signature pairs that belonged to the “multiple disease multiple tissue” category, a majority connected signatures issued from brain and blood, brain and islets and brain and vessel. Specifically, for schizophrenia - type 2 diabetes (scz-t2d) signature pairs, brain and islets, brain and adipose, and brain and liver were the major sources indicating again that there is no bias towards connecting signatures from the same tissue (Fig. S3). To summarize, the computed signature graph displayed numerous connected signature pairs across cohorts and tissues, with a majority of connections between signatures extracted from schizophrenia datasets and type 2 diabetes datasets.

**Figure 2.**
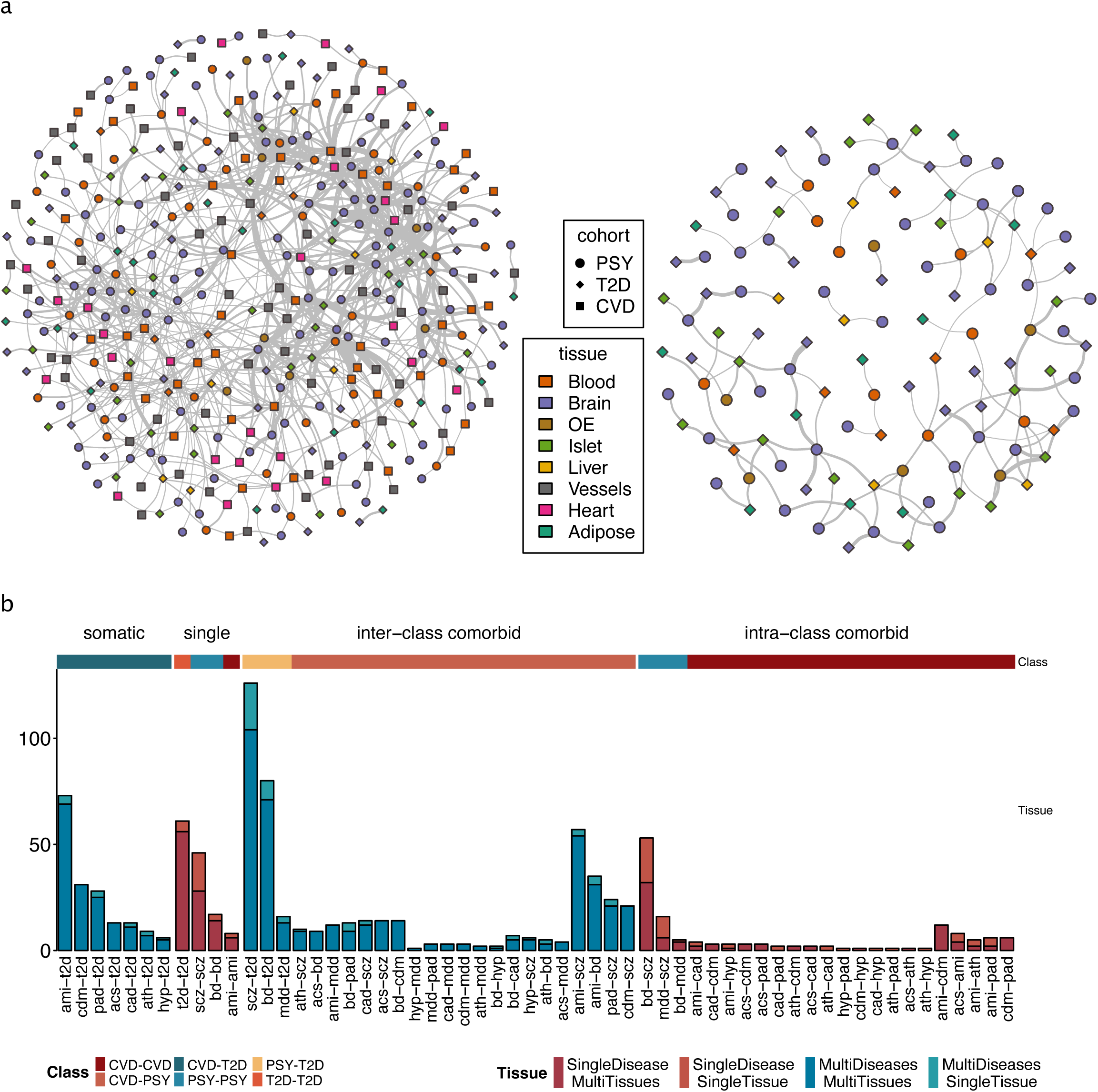
Computed signature graph and descriptive summary. **a left** Signature graph (full). Each node is an individual signature. Each edge indicates a reciprocal best hit matching between. Node shape represents disease class. Node color refers to the tissue origin. **right** Signature graph (scz-t2d). **b** Summary of the signature pairs (edges). y-values represent the number of the signature pairs in specific categories. Single category refers to cohorts that only contain signatures from one of the disease categories among PSY, CVD and T2D. Comorbid category consists of multiple signatures from PSY-CVD, PSY-T2D and PSY+CVD+T2D at the same time. Somatic category is composed of CVD+T2D signatures.

### Deciphering the variety of gene composition in comorbid signature pairs

Next, we wanted to understand to what extent the hybrid signatures connecting schizophrenia signatures to signatures from somatic diseases (called “comorbid pairs” in the remainder) contain specific biological signals that would not be identified by looking at each disease individually. To each pair of connected signatures, we associate the list of genes representing the intersection between the top 1000 genes in each one of the two signatures. We then performed a clustering based on the Jaccard distance between the gene lists associated with each pair of signatures. Several clusters were identified within and across signature pairs of scz-scz, t2d-t2d and scz-t2d respectively, suggesting that both distinct and specific sets of genes as well as biological modules were identified (Fig 3a). We also observed that some of comorbid signature pairs (e.g. scz-t2d, ami-scz) had a relatively high similarity with non-comorbid signature pairs (e.g. scz-scz, t2d-t2d, ami-ami). This observation suggested that part of the signal captured in the comorbid pairs might recapitulate processes already found in each single disease. To understand the amount of specific signal that can be found in the comorbid signature pairs, we started by exploring the genes in the gene list of comorbid signature pairs specifically. In some of the previous studies, comorbid genes between psychotic and its comorbidity were defined as the shared genes discovered by separate analyses on individual diseases ^6,9,10^. We reasoned that genes in the gene lists of signature pairs connecting identical diseases (i.e. scz-scz or t2d-t2d) represent the specific biological processes linked to the disease, while gene lists of comorbid pairs would represent the shared biological dimension (“comorbid genes”). Hence, we extracted the genes in scz-scz, t2d-t2d and scz-t2d signature pairs and compared them to previously identified gene sets. For instance, we labeled as *schizophrenia specific* the genes in our scz-t2d signature pairs that can only be found in the first single disease signature pair (e.g. scz-scz signature pairs), as *comorbidity specific* those that can only be found in the comorbid disease (e.g. t2d-t2d), as *shared* those that can be found in both scz-scz and t2d-t2d disease signature pair and *distinct* those that are not found in any of the single-disease signature pairs (Fig 3b). Looking at two comorbidities with schizophrenia, diabetes and acute myocardial infarction, we observe that 39% (scz-t2d) and 48% (scz-ami) of the comorbid genes are indeed distinct from all the other categories, that is they were not previously identified in the single-disease signature pairs (scz-scz or t2d-t2d and scz-scz or ami-ami) and represent potential novel comorbid genes (Fig 3b). These results indicate that the genes associated with comorbid signature pairs capture a specific biological signal that is distinct from the one explaining each disease separately.

**Figure 3.**
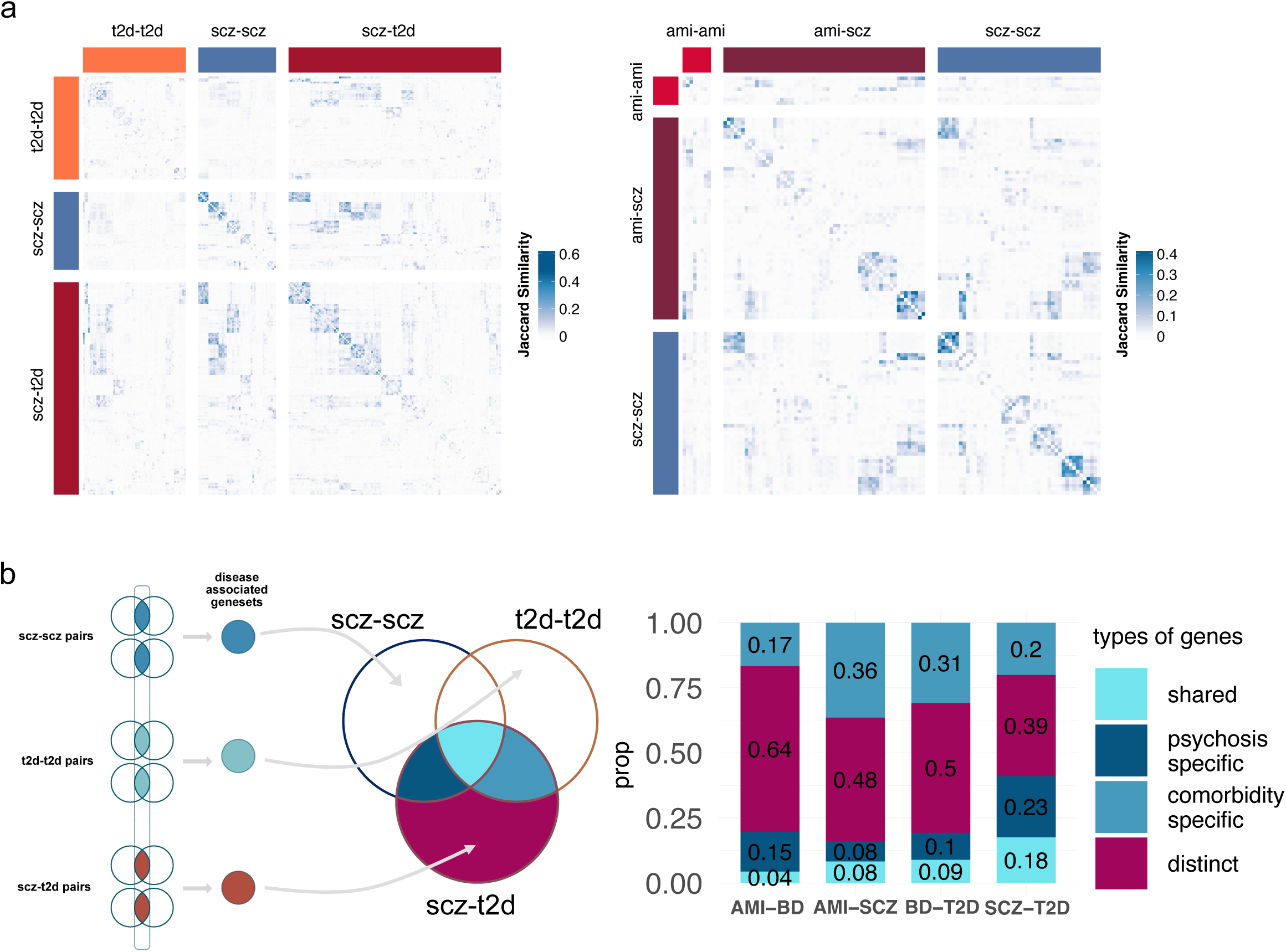
Identification of diverse types of comorbid signature pairs and the corresponding gene sets. **a** Jaccard similarity between scz-scz, scz-t2d and t2d-t2d signature pairs **b left** Illustration of categorizing types of genes in signature pairs, **right** Various types of comorbid genes in which the genes were labeled as “schizophrenia specific” in scz-t2d signature pairs that can only be found in the first single disease signature pair (e.g. scz-scz signature pairs), as “comorbidity specific” in scz-t2d signature pairs that can only be found in the first single disease signature pair (e.g. t2d-t2d or ami-ami signature pairs), as “shared” that can be found in the both scz-scz and t2d-t2d (or ami-ami) disease signature pair and “distinct” those that are not found in single disease signature pairs.

In summary, we revealed in the analysis that the set of comorbid genes includes both genes identified in single-disease analysis (i.e. in scz-scz and t2d-t2d disease signature pairs) as well as distinct genes found in neither of the two single diseases.

### Deciphering inflammatory response in schizophrenia comorbidity

In the remainder of this study, we decided to focus on the comorbidity between schizophrenia and diabetes. To explore the underlying mechanisms linked to comorbidity, we conducted a fine-grained functional enrichment analysis on each of the scz-t2d signature pairs using the overlapping gene lists as input (see Method). After clustering of similar terms, 27 clusters containing 590 enriched GO terms were identified. Many of these terms were found to be enriched across multiple signature pairs, which indicates the robustness of this enrichment. We identified for example numerous terms covering a wide range of activities of immune factors, such as the production, activation, migration and regulation of immune cells, and production of cytokines (Fig 4a). Interestingly, our analysis also identified an enrichment for *regulation of complement activation*, confirming the important role of the complement system as a shared biological dimension between both diseases. This analysis indicates that immunological processes and inflammatory responses might play an important role in the shared molecular dimension between schizophrenia and diabetes. This is consistent with numerous studies on possible comorbidity mechanisms, confirming the validity of our strategy ^12,27^. However, other shared biological processes were less expected. Within the cluster related to *anatomical structure development*, the processes related to *angiogenesis* and *vasculature development* were identified as being shared by 19 scz-t2d pairs. It has been described that regulation of the microvascular environment leads to vascular abnormalities inducing angiogenic factors and neuroinflammation in schizophrenia patients ^28–30^. Vascular abnormalities were also described in the pathogenesis of diabetes involving diabetic neuropathy such as changes of microvascular environment ^31,32^.

**Figure 4.**
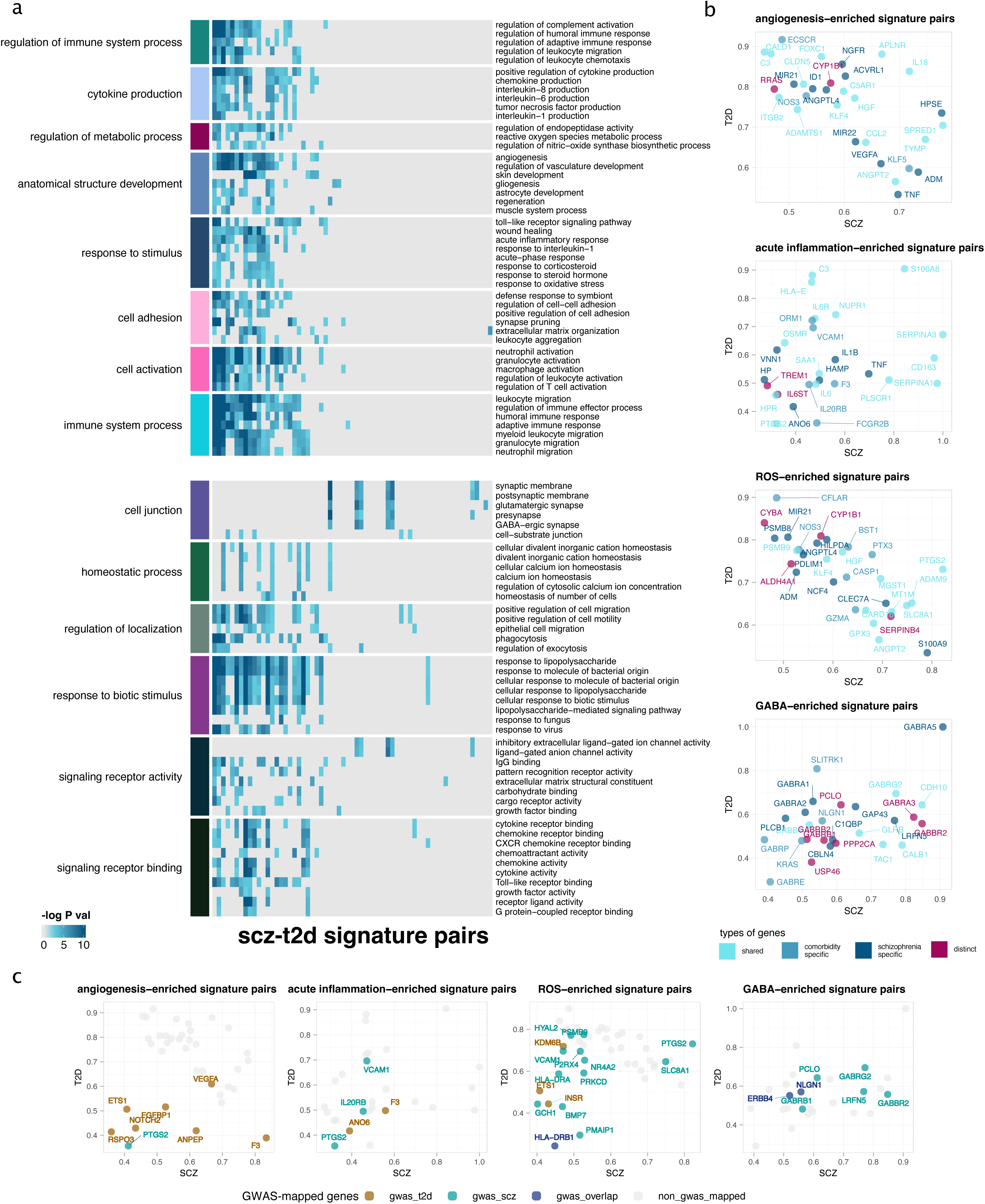
Biological representation of graph attributes. **a** Biological processes identified in scz-t2d signature pairs within clusters obtained by GO grouping on the enriched terms. Each column represented one individual signature pair. Left annotations are the cluster names while the rownames indicate the processes in each cluster. **b** Genes ranked by the exposure associated with the enriched pathways where x axis refers to the exposure of a gene in scz signature within a scz-t2d signature pair while y axis is the exposure of a gene in t2d signature within the signature pair. Colors refer to the type of genes which corresponds to the classification of various types of genes in Figure 3b. **C** Genes associated with GWAS risk variants

Interestingly, *wound healing process* was also identified. Indeed, impaired wound healing was described as one of the most disabling diabetic complications ^32^. In addition, a meta-analysis of various studies investigating the relationship between psychological stress and wound healing found a robust negative relationship between both ^33^.

Besides, *reactive oxygen metabolism* and *oxidase process* were also observed to be enriched in comorbid signature pairs. These impaired processes have been mentioned in several previous studies on t2d comorbidity of schizophrenia where oxidative-antioxidant imbalance led to the dysregulation of lipid transportation which caused and deteriorated the diabetes syndrome ^30,34^.

In addition, other processes also raised our interest though they were found in a smaller cluster. For instance, we observed one cluster annotated with synapse-related processes. In particular, *GABA-ergic synapse* was found to be enriched in 5 signature pairs. Furthermore, smaller clusters were also identified within other scz-t2d signature pairs involving, for example, cation homeostasis, immunoglobulin binding, and innate immune response.

To understand what specific genes were identified in the biological processes, we extracted the gene exposure from the signatures in the comorbid signature pairs (see Method, Fig 4b and Fig S1). We focussed on four processes: angiogenesis, acute inflammation, ROS and GABAergic processes. All the scz-t2d comorbid signature pairs enriched for these terms were retrieved and the top genes with the highest exposure were ranked (Fig 4b), including genes with top exposure in scz signatures and t2d signatures respectively and genes with top geometric mean exposure. The genes were labeled according to the categories they belong to (Fig. 3b). For genes related to angiogenesis-enriched signature pairs, we highlighted those with high exposure value in both scz and t2d signatures as well as the geometric mean exposure ranking, including *IL18, APLNR, NGFR, TYMP, SPRED1* and *HPSE*. Some of these genes including *IL18, APLNR, NGFR* and *SPRED1* were already shown to be dysregulated in various processes such as inflammation, energy metabolism, peripheral neuropathy and cell plasticity in both diseases ^35–43^. For instance, *APLNR,* encoding apelin APJ system is implicated in psychosis and neuropathy processes in chronic schizophrenia ^43,44^ but also acts as a promising biomarker for the treatment of type 2 diabetes given its functions in energy metabolism in insulin resistance progress via adipokines involvement ^45^. Other genes were not yet described in the context of comorbidities. Specifically, *TYMP,* which promotes angiogenesis and stimulates the growth of endothelial cells, was already found associated with endothelial dysfunction and induced diabetic-like symptoms in studies of type 2 diabetes ^46^, but has not been described in the context of schizophrenia. However, mitochondrial neurogastrointestinal dysfunction, associated with biallelic variants in the *TYMP* gene, was observed commonly in schizophrenia ^47,48^. We also investigated genes related to the enriched GABAergic system processes. Here, *GABRA5* was identified in scz-t2d comorbid genes. This gene is involved in neurotransmitter regulation in psychosis. In diabetic patients, the GABAergic system regulates GABA release in the endocrine system which induces membrane depolarization and increases insulin secretion ^49^. It is also associated with episodic memory dysfunction and lower cognitive performance ^50^. Other genes such as *CDH10,* encoding neuronal cell–adhesion molecules, were already shown to be associated with neuropsychiatric disorders (e.g. autism spectrum disorders)^51^ but not specifically to schizophrenia so far. In diabetes, *CDH10* and the cadherin family were identified in several complications of diabetes mellitus such as diabetic peripheral neuropathy^51,52^ and diabetic kidney disease ^51–53^.

To summarize, our analysis of comorbid signature pairs connecting schizophrenia and diabetes highlighted several enriched functional terms, among which acute inflammation, angiogenesis, oxidative process and GABAergic pathway. We noticed that most of the identified processes were to some extent associated with inflammatory response, therefore suggesting inflammation as a central mechanism in schizophrenia - type 2 diabetes comorbidity. Many of these processes had been connected separately to schizophrenia and diabetes in the literature, but our analysis strengthens the hypothesis that these processes represent shared molecular processes explaining the higher susceptibility to diabetes for schizophrenia patients.

### GWAS mapping of the embedded genes in the specific comorbid mechanism

To inspect the genes identified in the four specific comorbid mechanisms and strengthen their connection to the disease, we mapped GWAS variants for schizophrenia and diabetes onto the genes related to the identified processes (See Method). We retrieved the enriched genes in processes of acute inflammation, angiogenesis, oxidative process, and GABAergic systems. We classified the genes with GWAS variants in schizophrenia-related traits, in type 2 diabetes traits, and co-occurring in both traits into gwas_scz, gwas_t2d and gwas_shared genes. In total, 33 unique GWAS genes were mapped into the four specific comorbid processes, with oxidative-related processes matching most with 16 genes (Fig 4c). Some GWAS genes, like *PTGS2* and *F3* were found both in angiogenesis and acute inflammatory response. For angiogenesis, *VEGFA* is a risk variant specifically in t2d and has also been identified as an important biomarker regulating energy metabolism and vascular blood flow in schizophrenia ^54^. Gene with GWAS variants in both traits were found in oxidative-related processes and GABAergic systems. The neuregulin 1 gene (*NRGN1*) coupling with the receptor *ERBB4* was implicated in schizophrenia patients as one of the key dysregulated signaling in GABAergic circuits ^55^. Furthermore, human leukocyte antigen (HLA) molecules such as HLA-DRB1 trigger autoimmune reactions in schizophrenia ^56^ and diabetic ^57^ patients.

### Building the potential scz-t2d comorbidity mechanism through knowledge graphs

To summarize the potential mechanism underlying the comorbidity, a schizophrenia - type 2 diabetes comorbidity map was built using a disease knowledge graph specifically curated for schizophrenia, bipolar disorder and type 2 diabetes (See Method). We centered the analysis on the four processes (ROS, angiogenesis, inflammation and GABAergic processes), with a central focus on the inflammatory response, and identified the connecting path between the diseases, intending to illustrate the specific roles of the processes in connecting schizophrenia and type 2 diabetes. Finally, we identified the subgraphs with strong evidence for each process separately by extracting the paths connecting schizophrenia and type 2 diabetes (Figure 5).

**Figure 5.**
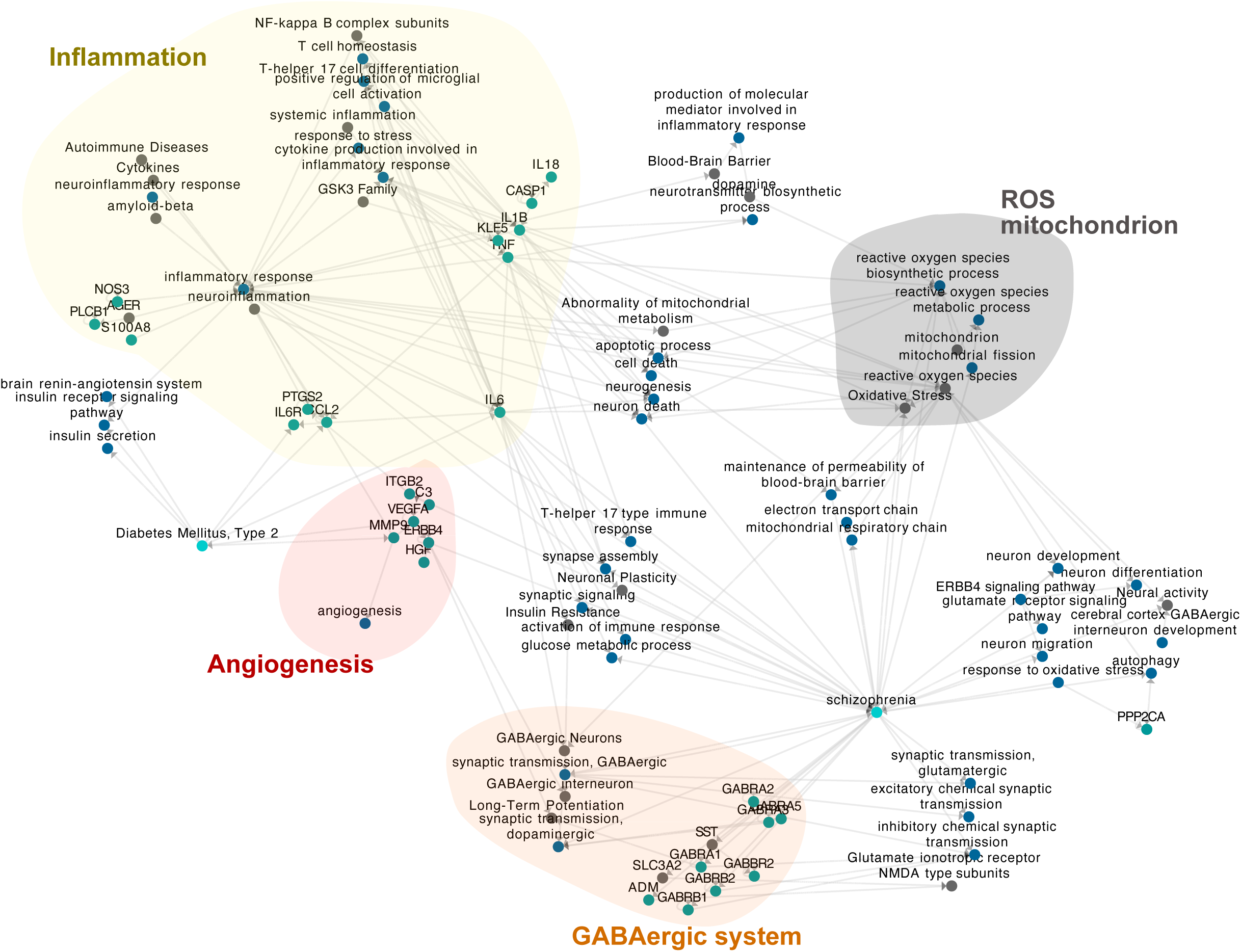
Schizophrenia - type 2 diabetes comorbidity knowledge-graph. Specific processes selected from all the previous analysis in the study including angiogenesis, inflammatory response, response to oxidative stress and GABA-ergic synapse, mapped on a knowledge-graph extracted from a literature-curated database. The output was generated and contained curated-connected nodes with maximal depth = 5 of these specific processes. Each node represents a specific entity (e.g. HGNC genes, GO biological processes, etc.) while each edge refers to a relationship or interaction between the connected nodes.

In this map, inflammation, as well as other related terms (neuroinflammation, systemic inflammation,…), appears as a highly connected hub node, with connections to the insulin receptor signaling pathway and hence to diabetes, as well as multiple connections, in particular through *IL6*, to schizophrenia. *IL6* and its receptor, *IL6R*, appear to directly connect diabetes to schizophrenia, with multiple connections bridging through the other highlighted processes including GABAergic neurons, oxidative stress, and neuronal plasticity. Molecular factors such as *TNF* and *IL6*, related to inflammatory response, have also been identified as top exposure genes in our previous analysis (Figure 4b).

Reactive oxygen species process played its roles via regulating the mitochondrial respiratory chain, electron transport chain, apoptotic process, in particular neuron death, and further processes related to the regulation of the blood-brain barrier, with multiple connections to schizophrenia. The connection to diabetes leads again through *IL6*. The angiogenesis term is connected to central genes such as *MMP9* and *VEGFA*, which establish the connection to the GABAergic system. Finally, terms in the knowledge graph related to the GABAergic system, as well as multiple associated genes are found to display numerous direct connections to schizophrenia, as well as to insulin resistance, confirming the role of GABAergic deregulation in the etiology of diabetes ^49^.

Based on the knowledge graph, we identified additional processes related to neural development (neuronal plasticity, neuronal signal transduction, neuron differentiation), energy metabolism (energy reserve metabolic process, mitochondrion, abnormality of mitochondrial metabolism), immune response (neuroinflammatory response, activation of immune response), blood-brain systems (blood-brain barrier, brain renin-angiotensin system), insulin system (insulin secretion, insulin receptor signaling) and cell death (apoptotic process, neuron death, autophagy). By combining these finding obtained from the knowledge graph and the enrichment analysis, we noticed that the co-occurrence of these processes forms a complex mechanism connecting not only oxidative response, cell death, but also other reactions such as cell self-protection, cell recovery, healing and regeneration, strengthening the hypothesis that the comorbidity of type 2 diabetes in schizophrenia could be inflammatory processes triggering damage and repair processes.

## Discussion

In summary, we presented a computational framework to perform comorbidity modeling via an improved integrative unsupervised machine learning approach based on multi-rank non-negative matrix factorization (mrNMF), based on the analysis of a large number of disease related transcriptomic datasets. Our approach addresses the parameter selection limitation of normal NMF by introducing multi-rank ensembled NMF to identify signatures in a hierarchical way. Using a robust network based approach, we have identified modules of connected disease signatures. A large number of these connections related signatures from distinct diseases, highlighting the shared transcriptomic dimension between different diseases. We have focused our analysis on the relationship between schizophrenia and diabetes, and found significant enrichment in shared transcriptomic signatures related to several processes: inflammatory processes, reactive-oxygen related processes, processes related to cell motility and angiogenesis as well as processes related to the GABAergic system. The relevance of these processes was confirmed in an independent analysis of a literature-derived knowledge graph.

The presented workflow can be extended to other cross-cohort comorbidity investigations, using reciprocal best-hit scoring matrices to control the batch effects. This computational approach can be adapted to numerous other biological problems. For instance, various types of data could be applied with the framework such as proteomics and metabolomics, extensively, cross modality data (e.g. genomics and proteomics) could also be integrated simultaneously using the workflow. Research purposes other than comorbidity investigation are also applicable. For example, transcription factor network analysis in either the same disease or multiple diseases are possible. Drug-drug combination networks inferred from gene expression profiles can also be studied.

Of note, several limitations in our analysis should be put into consideration. The overlapping pathways were identified when disease-associated pathways co-occurred in both diseases, when we curated our reference gene set. However, we cannot always distinguish between causative effects, leading to real comorbidity risk, and shared downstream consequences of each disease. However, our GWAS analysis appears to confirm several genes identified by our unsupervised analysis as shared risk variants in diabetes and schizophrenia, such as *ERBB4* and *NLGN1*. *NLGN1* is an interesting hit, for its role in the interaction of endothelial cells with extracellular matrix during angiogenesis and its downregulation in diabetic patients^58^. At the same time, variants of the paralogous *NLGN2* gene have been identified as risk variants in schizophrenia due to their role as cell-adhesion molecules in post-synaptic membrane ^59^. This variant is associated with loss-of-function in the formation of GABAergic synapses, connecting back to one of the key processes identified in our analysis.

Overall, the computational framework based on mrNMF signature graph is a powerful and unbiased approach for cross-cohort modeling of disease comorbidity. By means of ensembling multi-ranks, it captures transcriptomic signatures at different levels of granularity, providing a complete description of cellular processes that might underlie disease comorbidities.

## Acknowledgements

YZ and CH are supported by the e:Med project COMMITMENT (Grant 01ZX1904D) from the German ministry for research and education (BMBF). We thank Emanuel Schwarz for discussions on this work.

---

**Supplementary Table S1.** Summary of datasets used in analysis.

|  | dataset | numsa<br>mple | numco<br>ntrol | num<br>case | cond<br>ition | cohort | tissue | platform | reference |  |  |  |  |  |  |  |  |  |  |
| --- | --- | --- | --- | --- | --- | --- | --- | --- | --- | --- | --- | --- | --- | --- | --- | --- | --- | --- | --- |
| 1 | GSE21138 | 59 | 29 | 30 | SCZ | PSY | Brain | GPL570[HG-U133_Plus_2]<br>Affymetrix Human<br>Genome U133 Plus 2.0<br>Array | https://www.ncbi.nlm.nih.gov/geo/query/acc.cgi?acc=GSE21138 | 14 | GSE15932 | 16 | 8 | 8 | T2D | T2D | Blood | GPL570[HG-U133_Plus_2]<br>Affymetrix Human<br>Genome U133 Plus 2.0<br>Array | https://www.ncbi.nlm.nih.gov/geo/query/acc.cgi?acc=GSE15932 |
| 2 | GSE17612 | 49 | 22 | 27 | SCZ | PSY | Brain | GPL570[HG-U133_Plus_2]<br>Affymetrix Human<br>Genome U133 Plus 2.0<br>Array | https://www.ncbi.nlm.nih.gov/geo/query/acc.cgi?acc=GSE17612 | 15 | GSE71416 | 20 | 6 | 14 | T2D | T2D | Adipose | GPL570[HG-U133_Plus_2]<br>Affymetrix Human<br>Genome U133 Plus 2.0<br>Array | https://www.ncbi.nlm.nih.gov/geo/query/acc.cgi?acc=GSE71416 |
| 3 | GSE27383 | 51 | 29 | 22 | SCZ | PSY | Blood | GPL570[HG-U133_Plus_2]<br>Affymetrix Human<br>Genome U133 Plus 2.0<br>Array | https://www.ncbi.nlm.nih.gov/geo/query/acc.cgi?acc=GSE27383 | 16 | GSE38396 | 8 | 4 | 4 | T2D | T2D | Skin | GPL570[HG-U133_Plus_2]<br>Affymetrix Human<br>Genome U133 Plus 2.0<br>Array | https://www.ncbi.nlm.nih.gov/geo/query/acc.cgi?acc=GSE38396 |
| 4 | GSE53987_2 | 70 | 18 | 52 | BD | PSY | Brain | GPL570[HG-U133_Plus_2]<br>Affymetrix Human<br>Genome U133 Plus 2.0<br>Array | https://www.ncbi.nlm.nih.gov/geo/query/acc.cgi?acc=GSE53987 | 17 | GSE24752 | 6 | 3 | 3 | HYP | CVD | Blood | GPL570[HG-U133_Plus_2]<br>Affymetrix Human<br>Genome U133 Plus 2.0<br>Array | https://www.ncbi.nlm.nih.gov/geo/query/acc.cgi?acc=GSE24752 |
| 5 | GSE53987_3 | 69 | 19 | 50 | MDD | PSY | Brain | GPL570[HG-U133_Plus_2]<br>Affymetrix Human<br>Genome U133 Plus 2.0<br>Array | https://www.ncbi.nlm.nih.gov/geo/query/acc.cgi?acc=GSE53987 | 18 | GSE19339 | 8 | 4 | 4 | ACS | CVD | Vessels | GPL570[HG-U133_Plus_2]<br>Affymetrix Human<br>Genome U133 Plus 2.0<br>Array | https://www.ncbi.nlm.nih.gov/geo/query/acc.cgi?acc=GSE19339 |
| 6 | GSE74358 | 28 | 14 | 14 | BD | PSY | Brain | GPL570[HG-U133_Plus_2]<br>Affymetrix Human<br>Genome U133 Plus 2.0<br>Array | https://www.ncbi.nlm.nih.gov/geo/query/acc.cgi?acc=GSE74358 | 19 | GSE48060 | 52 | 21 | 31 | AMI | CVD | Blood | GPL570[HG-U133_Plus_2]<br>Affymetrix Human<br>Genome U133 Plus 2.0<br>Array | https://www.ncbi.nlm.nih.gov/geo/query/acc.cgi?acc=GSE48060 |
| 7 | GSE46449 | 88 | 39 | 49 | BD | PSY | Blood | GPL570[HG-U133_Plus_2]<br>Affymetrix Human<br>Genome U133 Plus 2.0<br>Array | https://www.ncbi.nlm.nih.gov/geo/query/acc.cgi?acc=GSE46449 | 20 | GSE66360 | 99 | 50 | 49 | AMI | CVD | Vessels | GPL570[HG-U133_Plus_2]<br>Affymetrix Human<br>Genome U133 Plus 2.0<br>Array | https://www.ncbi.nlm.nih.gov/geo/query/acc.cgi?acc=GSE66360 |
| 8 | GSE73129 | 115 | 60 | 55 | SCZ | PSY | OE | GPL570[HG-U133_Plus_2]<br>Affymetrix Human<br>Genome U133 Plus 2.0<br>Array | https://www.ncbi.nlm.nih.gov/geo/query/acc.cgi?acc=GSE73129 | 21 | GSE13985 | 10 | 5 | 5 | ATH | CVD | Blood | GPL570[HG-U133_Plus_2]<br>Affymetrix Human<br>Genome U133 Plus 2.0<br>Array | https://www.ncbi.nlm.nih.gov/geo/query/acc.cgi?acc=GSE13985 |
| 9 | GSE7036 | 6 | 3 | 3 | BD | PSY | Brain | GPL570[HG-U133_Plus_2]<br>Affymetrix Human<br>Genome U133 Plus 2.0<br>Array | https://www.ncbi.nlm.nih.gov/geo/query/acc.cgi?acc=GSE7036 | 22 | GSE97320 | 6 | 3 | 3 | AMI | CVD | Blood | GPL570[HG-U133_Plus_2]<br>Affymetrix Human<br>Genome U133 Plus 2.0<br>Array | https://www.ncbi.nlm.nih.gov/geo/query/acc.cgi?acc=GSE97320 |
| 10 | GSE21935 | 42 | 19 | 23 | SCZ | PSY | Brain | GPL570[HG-U133_Plus_2]<br>Affymetrix Human<br>Genome U133 Plus 2.0<br>Array | https://www.ncbi.nlm.nih.gov/geo/query/acc.cgi?acc=GSE21935 | 23 | GSE71226 | 6 | 3 | 3 | CAD | CVD | Blood | GPL570[HG-U133_Plus_2]<br>Affymetrix Human<br>Genome U133 Plus 2.0<br>Array | https://www.ncbi.nlm.nih.gov/geo/query/acc.cgi?acc=GSE71226 |
| 11 | GSE76894 | 97 | 78 | 19 | T2D | T2D | Islet | GPL570[HG-U133_Plus_2]<br>Affymetrix Human<br>Genome U133 Plus 2.0<br>Array | https://www.ncbi.nlm.nih.gov/geo/query/acc.cgi?acc=GSE76894 | 24 | GSE19303 | 48 | 8 | 40 | CDM | CVD | Heart | GPL570[HG-U133_Plus_2]<br>Affymetrix Human<br>Genome U133 Plus 2.0<br>Array | https://www.ncbi.nlm.nih.gov/geo/query/acc.cgi?acc=GSE19303 |
| 12 | GSE76895 | 57 | 27 | 30 | T2D | T2D | Islet | GPL570[HG-U133_Plus_2]<br>Affymetrix Human<br>Genome U133 Plus 2.0<br>Array | https://www.ncbi.nlm.nih.gov/geo/query/acc.cgi?acc=GSE76895 | 25 | GSE53987_1 | 66 | 18 | 48 | SCZ | PSY | Brain | GPL570[HG-U133_Plus_2]<br>Affymetrix Human<br>Genome U133 Plus 2.0<br>Array | https://www.ncbi.nlm.nih.gov/geo/query/acc.cgi?acc=GSE53987 |
| 13 | GSE23343 | 17 | 7 | 10 | T2D | T2D | Liver | GPL570[HG-U133_Plus_2]<br>Affymetrix Human<br>Genome U133 Plus 2.0<br>Array | https://www.ncbi.nlm.nih.gov/geo/query/acc.cgi?acc=GSE23343 | 26 | GSE161355 | 33 | 15 | 18 | T2D | T2D | Brain | GPL570[HG-U133_Plus_2]<br>Affymetrix Human<br>Genome U133 Plus 2.0<br>Array | https://www.ncbi.nlm.nih.gov/geo/query/acc.cgi?acc=GSE161355 |
|  |  |  |  |  |  |  |  |  |  | 27 | GSE27034 | 37 | 18 | 19 | PAD | CVD | Blood | GPL570[HG-U133_Plus_2]<br>Affymetrix Human<br>Genome U133 Plus 2.0<br>Array | https://www.ncbi.nlm.nih.gov/geo/query/acc.cgi?acc=GSE27034 |

**Supplementary Figure S1.**
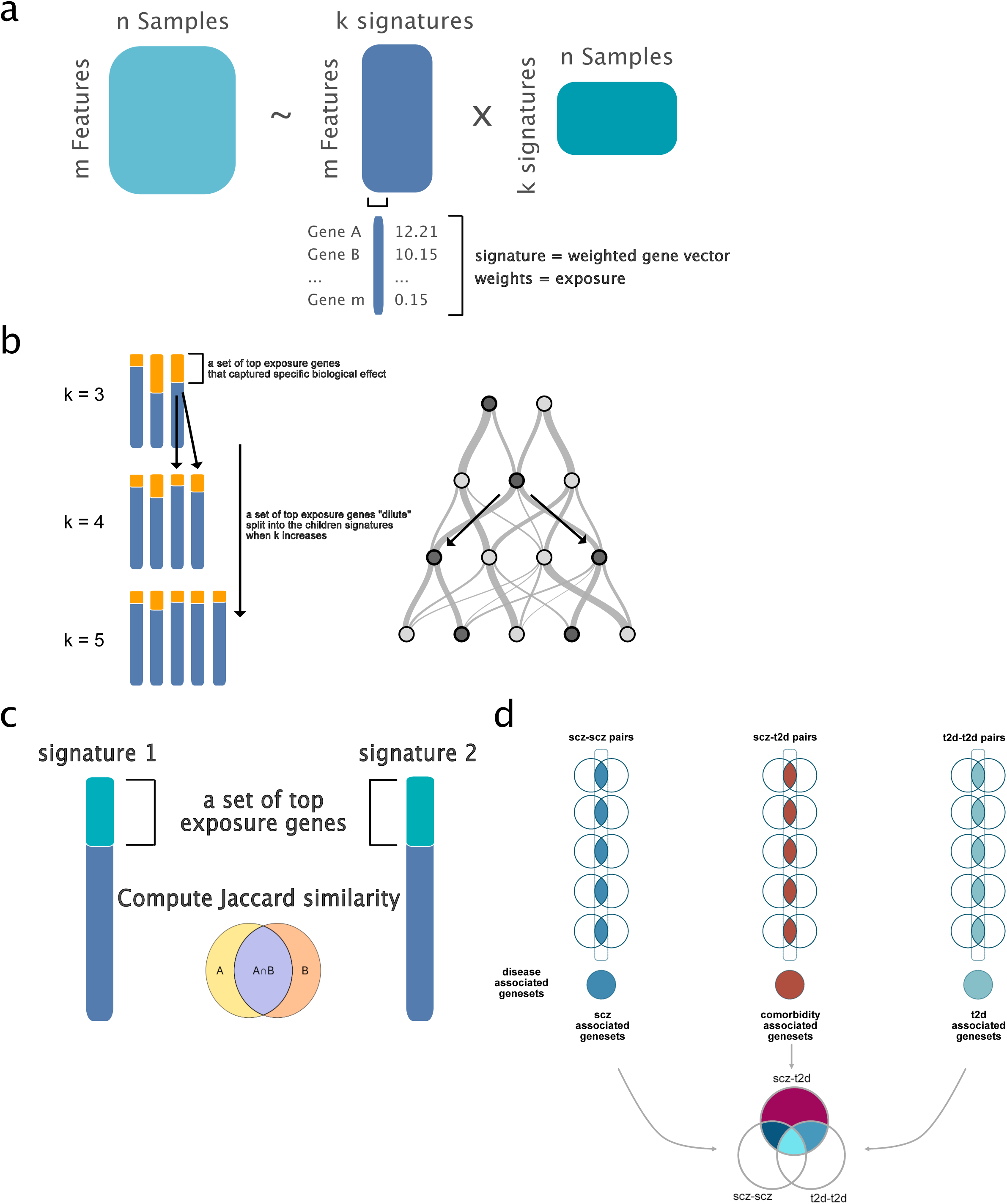
**a** Illustration of the non-negative matrix factorization and the matrix outputs **b left** Matrix decomposition with a series of different rank parameters k from k = 2 to k = 20 **right** Riverplot of the matrix factorization with a series of different rank where each node refers to signature, width of the flow refers to the similarity of the signatures and colors were annotated to indicate the diagnosis association of the signatures **c** Jaccard similarity calculated by selected top N of the genes from the signatures **d** Extraction of disease-associated genesets and compared across different genesets to identify various types of genes

**Supplementary Figure S2.**
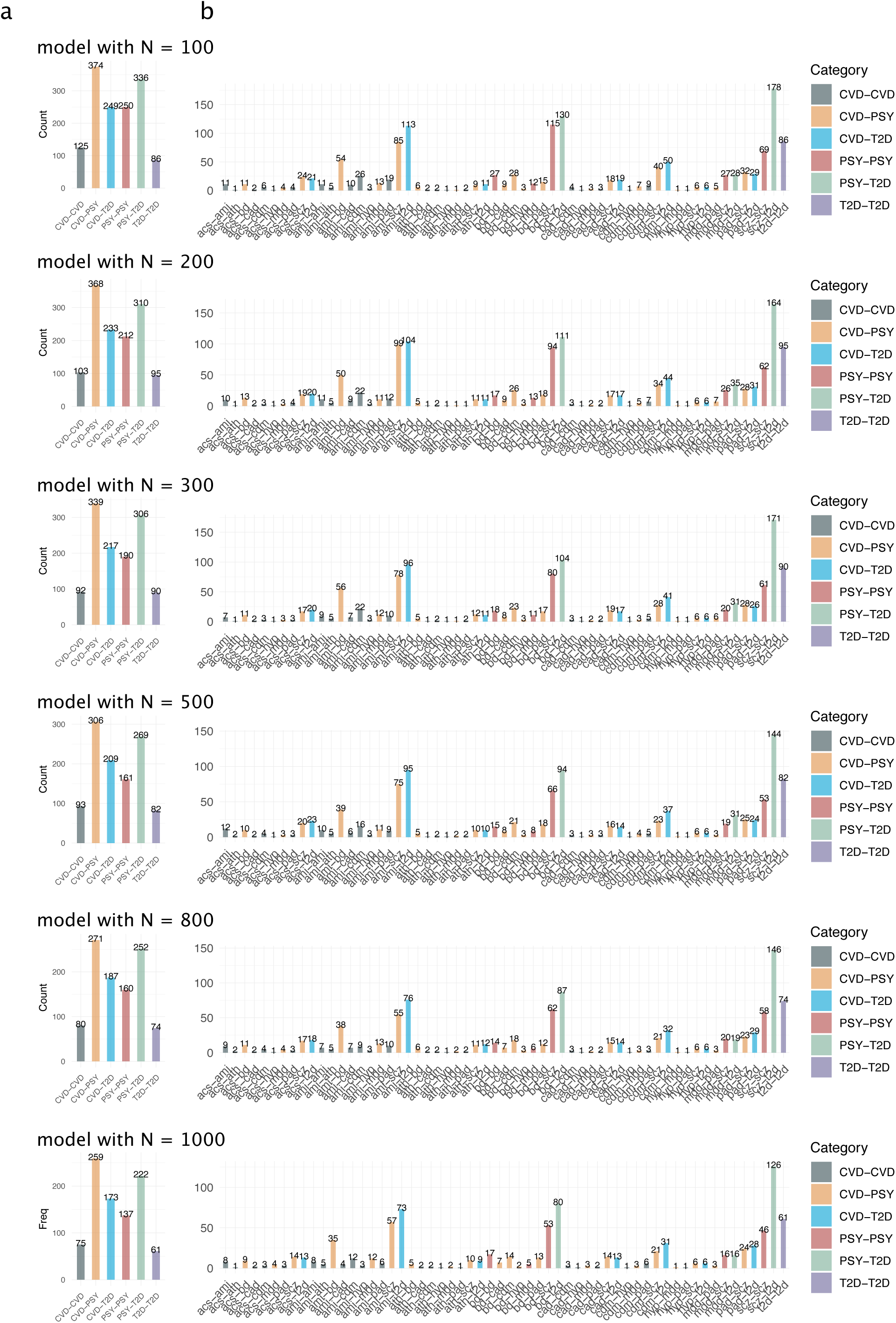
Summary of the identified signature pairs based on models with different top N genes. **a** Counts of identified signature pairs based on different models by the disease major class (e.g. PSY, T2D and CVD) **b** Counts of identified signature pairs based on different models by the condition class (e.g. scz, t2d, ami, etc..)

**Supplementary Figure S3.**
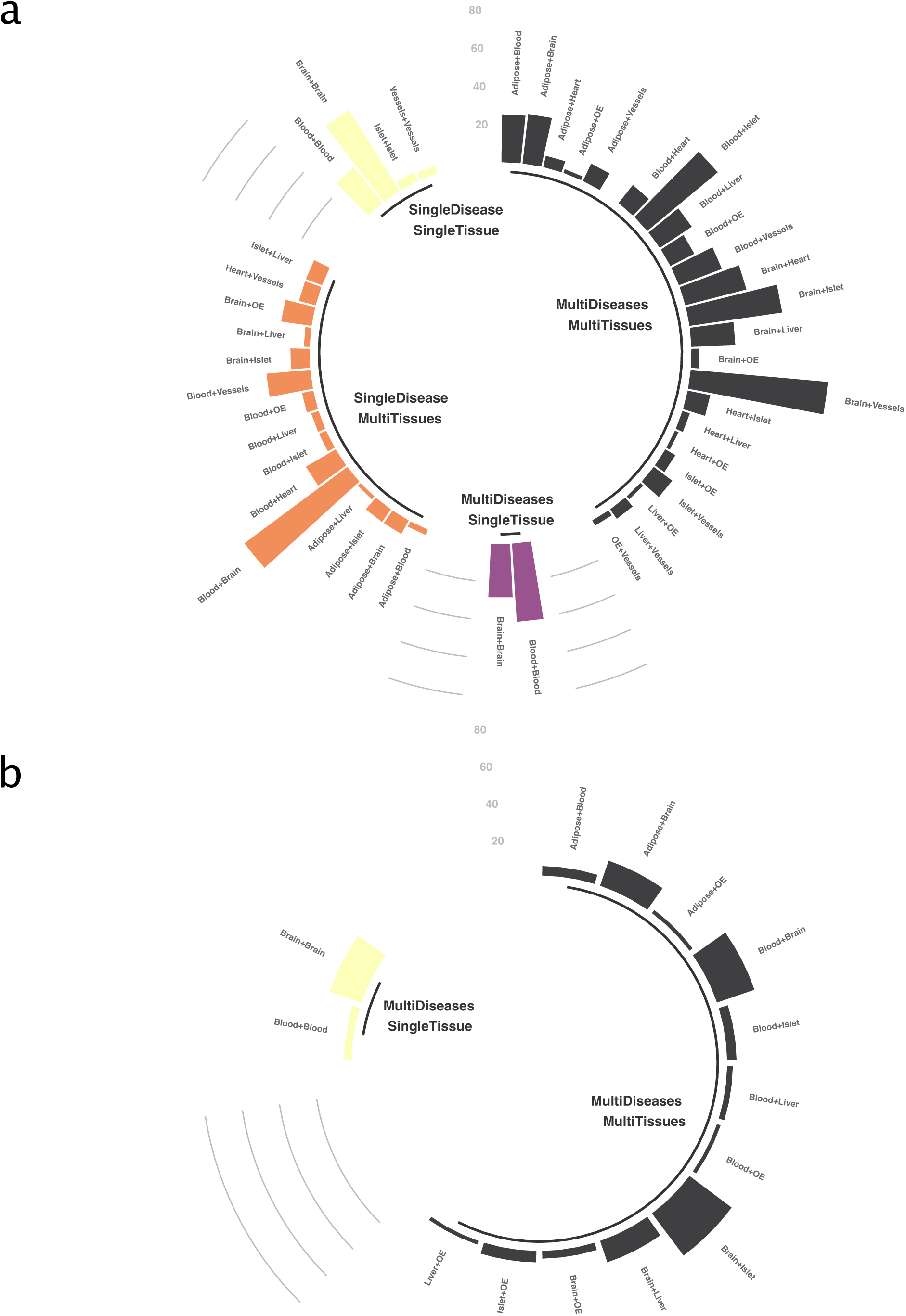
**a** Summary of the tissue-tissue combination of all the signature pairs **b** Summary of the tissue-tissue combination of the schizophrenia - type 2 diabetes signature pairs

